# Association of common genetic variants in the *CPSF7* and *SDHAF2* genes with Canine Idiopathic Pulmonary Fibrosis in the West Highland White Terrier

**DOI:** 10.1101/2020.04.14.030486

**Authors:** Ignazio S. Piras, Christiane Bleul, Ashley Siniard, Amanda J. Wolfe, Matthew D. De Both, Alvaro G. Hernandez, Matthew J. Huentelman

## Abstract

Canine Idiopathic Pulmonary Fibrosis (CIPF) is a chronic fibrotic lung disease that is observed at a higher frequency in the West Highland White Terrier dog breed (WHWT) and may have molecular pathological overlap with human lung fibrotic disease. We conducted a Genome-Wide Association Study (GWAS) in the WHWT using Whole Genome Sequencing (WGS) to discover genetic variants associated with CIPF. Saliva-derived DNA samples were sequenced using the Riptide™ DNA library prep kit. After quality controls, 28 affected, 44 unaffected and 1,843,695 informative Single Nucleotide Polymorphisms (SNPs) were included in the GWAS. Data were analyzed both at the single SNP and gene levels using the GEMMA and GATES methods, respectively. We detected significant signals at the gene level in both the *CPSF7* and *SDHAF2* genes (adjusted p = 0.016 and p = 0.025, respectively), two overlapping genes located on chromosome 18. The top SNP for both genes was rs22669389, however it did not reach genome-wide significance in the GWAS (adjusted p = 0.078). Our studies provide, for the first time, candidate loci for CIPF in the WHWT. *CPSF7* was recently associated with lung adenocarcinoma further highlighting the potential relevance of our results since IPF and lung cancer share several pathological mechanisms.

## 1. Introduction

Canine Idiopathic Pulmonary Fibrosis (CIPF) is a chronic and progressive fibrotic lung disease that particularly affects the West Highland White Terrier dog breed (WHWT) [1]. CIPF shares several clinical and pathological features with human IPF and it has been proposed as a possible sporadic disease model [2]. A typical clinical feature occurring in the majority of affected dogs is inspiratory crackles, as well as laryngo-tracheal reflex, tachypnea, and excessive abdominal breathing [2,3]. Human IPF is considered to be a disease of the epithelium. Specifically, microscopic injuries of the aging lung epithelium leads to defective regeneration and abnormal epithelial-mesenchymal crosstalk with activation of transforming growth factor (TGF-β) [4,5]. This is followed by extravascular coagulation, immune system activation, and secretion of excessive amounts of extracellular matrix (ECM) proteins [2]. However, the overall initiating cause of this pathological cascade in dogs and humans is still unknown.

Several studies have attempted to clarify the molecular mechanisms of CIPF in WHWTs. Maula et al. [6] reported the upregulation of Activin A in the lung alveolar epitelium of WHWTs with CIPF. Furthermore, increased TGF-β1 signaling activity was detected in WHWTs and other predisposed breeds (such as Scottish Terriers and Bichons Frise) compared to non-predisposed breeds [7]. In an RNA expression profiling study in dogs, chemokine and interleukin genes were found to be overexpressed in the lungs with CCL2 levels also noted to be elevated in serum [8]. Endothelin-1, measured in both serum and bronchoalveolar lavage fluid, was suggested to be a biomarker suitable to differentiate dogs with CIPF from dogs with chronic bronchitis and eosinophilic bronchopneumopathy [9].

Genetic background is considered one of the risk factors for both CIPF and IPF [3,10]. Genome-Wide association studies (GWAS) have led to the detection of several genes linked with the disease in humans. Specifically, three GWAS have been conducted in humans detecting signals in AKAP13, MUC5B, DSP, TOLLIP, MDGA2, SPPL2C and TERT [10–12]. However, genetic risk factors for CIPF in the WHWT have not been identified. The domestic dog (Canis familiaris) is a useful model for many human diseases [13] due to the high number of analogous diseases [14], similar physiologies and medical care, as well as the simplified genetic architecture in purebred dogs [15]. Each dog breed originated from a small founder population with consequently low levels of genomic heterogeneity and long stretches of linkage disequilibrium (LD). Due to these characteristics, GWAS in dogs have increased statistical power comparable to, or better than, those performed in human population isolates [16]. This genetic homogeneity, the consequence of strong artificial selection conducted by humans, also led to an excess of inherited diseases, offering unique opportunities to discover genetic associations for spontaneous disease [13]. Several GWAS have been conducted in specific breeds using the canine SNP array leading to the discovery of genetic risk factors for ectopic ureters [17], inflammatory bowel disease [18], hereditary ataxia [19], and hypothyroidism [20], among others.

In this study we conducted, for the first time, a GWAS in a sample WHWTs with CIPF and unaffected controls using whole genome sequencing and imputation with the goal of finding genetic risk factors that may predispose the WHWT to the disease.

## 2. Methods

### 2.1. Sample collection

A total of 73 dogs, including 28 affected (AF) and 45 (UF) were sampled via internet-based recruitment of saliva samples (the study site can be found at www.tgen.org/westie). Participants were directed to self-report the breed and diagnostic status of their dog. The complete enrollment form can be found in File S1. The study protocol was reviewed and approved by TGen’s Institutional Animal Care and Use Committee (Protocols #13055, #15002, #18006) in accordance with relevant guidelines and regulations. Owner consent for collection of the samples used in this study was obtained. The sex ratio (male:females) in AF was 0.86, whereas in UF was 1.26 (*p* = 0.466). The average age for AF was 12.8 years (range: 6.9 - 17.0), whereas in UF was 12.7 years (range: 9.1 – 16.8) (*p* = 0.793). Sex information was not available for 4 samples, age information was not available for one sample. Saliva was collected by the owner using the Oragene ANIMAL kit (DNAGenotek, Ottawa, CA) and returned to the laboratory at ambient temperature.

### 2.2. DNA extraction and whole genome sequencing

DNA was isolated from the collected saliva specimens according to the manufacturer’s instructions. Construction of the shotgun genomic libraries and sequencing on the NovaSeq 6000 was carried out at the Roy J. Carver Biotechnology Center, University of Illinois at Urbana-Champaign. DNA was quantitated with the Qubit High Sensitivity reagent (Thermo Fisher, Waltham, MA) and diluted with water to 2.5 ng/μl in a total volume of 12ul. Libraries were prepared with the Riptide™ DNA library prep kit (iGenomX, Carlsbad, CA). Briefly, random primers with 5’ barcoded Illumina adapter sequences (one sequence that is unique to each sample) are annealed to denatured DNA template. A polymerase extends each primer and this action is terminated with a biotinylated dideoxynucleotide, of which there is a small fraction in the nucleotide mix. The biotinylated products are then pooled for all of the samples and captured on streptavidin coated magnetic beads. A second 5’adapter-tailed random primer is used with a strand-displacing polymerase to convert the captured DNA strands into a dual adapter library. PCR is used to amplify the products and add an index barcode. These libraries were then sequenced for 150nt from each side of the DNA fragments (paired-reads) on a NovaSeq 6000 (Illumina, San Diego, CA) one lane of an S2 flowcell.

### 2.3. Data analysis

The fastq files were generated with the bcl2fastq v2.20 Conversion Software (Illumina) and demultiplexed with the fgbio tool from Fulcrum Genomics (https://github.com/fulcrumgenomics/fgbio). Sequencing reads were processed and imputed using version 2.0 of the Gencove, Inc. analysis pipeline for canine low-pass sequencing data. Reads were aligned to the reference genome canFam3.1 using bwa mem v0.7.17, sorted and duplicates marked using samtools v1.8, and imputation performed using loimpute v0.18 (Gencove, Inc.), based on the model of Li et al. [21] The imputation reference panel consists of 676 sequenced dogs across the 91 dog breeds for a total of 53 Millions of sites.

The resulting *vcf* files of 28 AF and 45 UF were filtered using *vcftools v0.1.16* including only biallelic, single nucleotide variants (SNVs) and variants with genotype probability (GP, indicating the imputation quality) greater or equal then 0.90. Then, we filtered the dataset with *PLINK v1.9* using the following thresholds: SNP genotyping rate ≥ 95%, MAF ≥ 5%, Hardy-Weinberg equilibrium in unaffected *p* ≥ 1.0E-05, sample genotyping rate ≥ 90%, and keeping only autosomal variants. We conducted principal component analysis (PCA) with *PLINK v1.9* to detect and remove outliers. Specifically, we used the identity by similarity (IBS) metric (taking into account from the 1^st^ to the 5^th^ closest neighbor, and classifying as outliers samples with Z ≤ −4, representing 4 standard deviations below the group mean. After outlier removal, the original dataset including only high-quality imputed SNPs was filtered again with *PLINK v1.9* using the same thresholds. Identity by Descent (IBD) analysis was conducted to estimate the relatedness between all the pair of samples calculating the *pi-hat* value using the --*genome* command in *PLINK v1.9*.

The GWAS was conducted using a Mixed Linear Model (MLM) to account for relatedness and population structure, as implemented in the *GEMMA v0.96* software [28], assessing the significance with the Wald test. Results were corrected at the genome-wide level using the Bonferroni method, accounting for the number of independent SNPs tested according the Linkage Disequilibrium patterns estimated using the option --*indep-pairwise 10000 1 0.80* in *PLINK v1.9*. Using this approach, we found 101,740 independent SNPs. Variants were annotated according *CanFam3.1* assembly using the R-package *Biomart v2.42.0*. Lambda inflation factor (*λ*) and quantile-quantile plots (qqplots) were computed using the R-package *snpStats*.

We further analyzed the data conducting a gene-based association analysis, using the GATES method [24], using as input the summary statistics obtained from the GEMMA analysis. First, we filtered the dataset including the SNPs located at ± 1,500 bp from each gene, to include the variants located in the promoter and in the 3’ regions. The analysis was also conducted including larger regions (± 5,000 and ± 10,000 bp). Ensemble start and end coordinates of each gene were retrieved using the R-packaged *Biomart v2.42.0*, according the dog assembly *CanFam3.1*. The Ensembl Biomart gene coordinates correspond to the outermost transcript start and end. Then, for each gene we computed a matrix of correlation between SNPs using the unaffected samples, in order to account for the LD. The correlation matrix was computed with the Pearson method using the *cor* function implemented in R, using the option *use = “na.or.complete”* to deal with missing genotypes. Finally, the *GATES* statistics were computed using as input the *p* values from the MLM analysis and the correlation LD matrix. The analysis was conducted using the *GATES2* function as implemented in the R-package *aSPU* (https://cran.r-project.org/web/packages/aSPU/index.html). P values were corrected using the Bonferroni method adjusting for the total number of genes tested.

## 3. Results

### 3.1. Filtering and quality controls

After imputation we removed INDELS, low quality variants, and variants with more than 2 alleles. We obtained a median number of SNVs per sample equal to 35,916,311 (range: 33,918,432 – 36,243,510). Samples showed a median depth of 1X. The median value of variants covered with equal to or greater than five reads per sample was 1,902,261 (range: 34,490 −12,200,995) (Table S1). We filtered the dataset with *PLINK v1.9* obtaining 1,839,683 variants for study with an average genotyping rate equal to 98.0%. We conducted PCA and IBS analyses to identify significant outliers and identified one outlier, based on the first and second principal components, with a Z score < −4 (Figure S1). We removed this sample from the analysis and we filtered the dataset again obtaining 1,843,695 SNPs in 28 AF and 44 UF animals. We re-ran the PCA and IBS analyses and did not identify any additional outliers. AF and UF groups were not statistically different for sex (p = 0.466) or age (*p* = 0.793). The IBD analysis demonstrated a *pi-hat* = 0.043 ± 0.068 (Range: 0.000 – 0.500). The distribution of *pi-hat* values for each sample pair is reported in Figure S2.

### 3.2. Genome-wide association study

We ran the GWAS accounting for relatedness and population stratification using the linear mixed model as implemented in the *GEMMA* software [26]. Age and sex were not included as covariates in the model because these factors did not differ significantly between the AF and UF animals. The results were adjusted with the Bonferroni method accounting for the 101,740 independent SNPs estimated by regional LD patterns, setting the genome-wide significance threshold at *p* < 4.91E-07 (adj *p* < 0.05). We considered a level of *p* < 9.83E-07 (adj *p* < 0.10) as “suggestive” association. The top 10 variants ranked by adj p value are reported in Table 1 and the Manhattan plot with the top 500K SNPs is illustrated in Figure 1A.

**Table 1.**
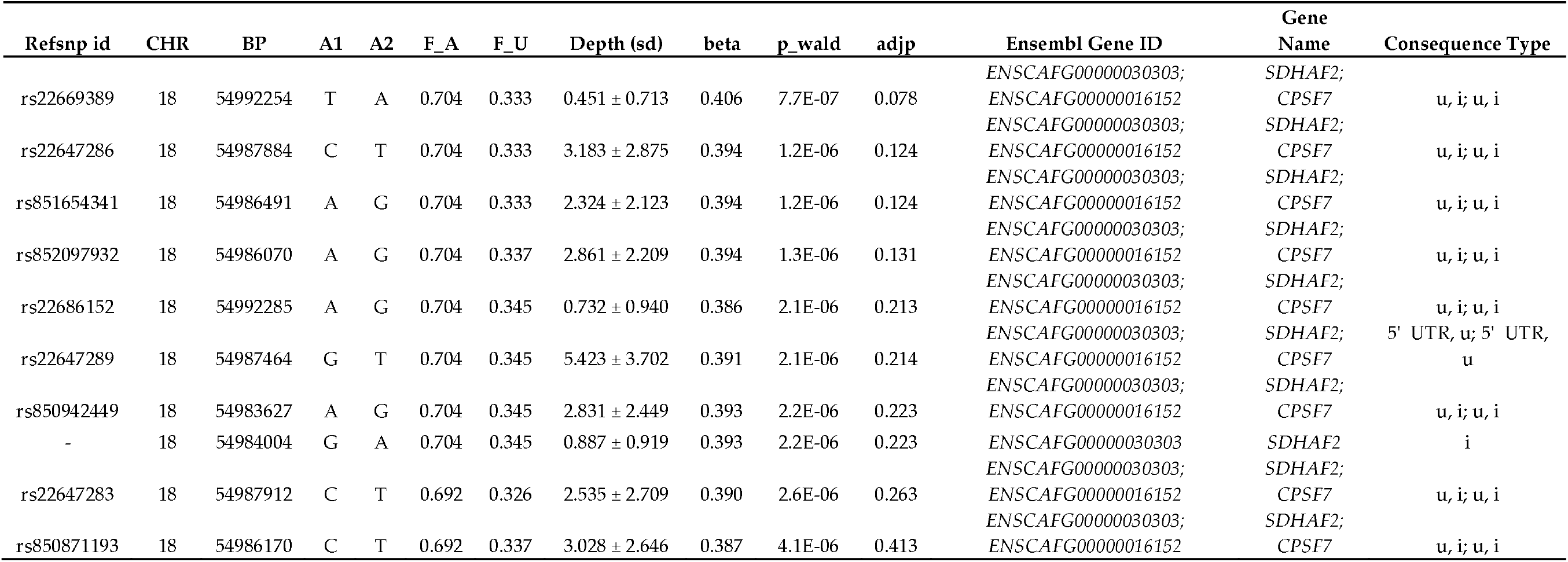
Details of the top 10 SNPs detected in the GWAS. p-values were adjusted using the Bonferroni method accounting for 101,740 independent SNPs A1: minor frequency allele referred to the total sample; A2: major frequency allele referred to the total sample; AF: frequency of A1 in affected; UF: frequency of A2 in unaffected; u: upstream; i: intron.

**Figure 1.**
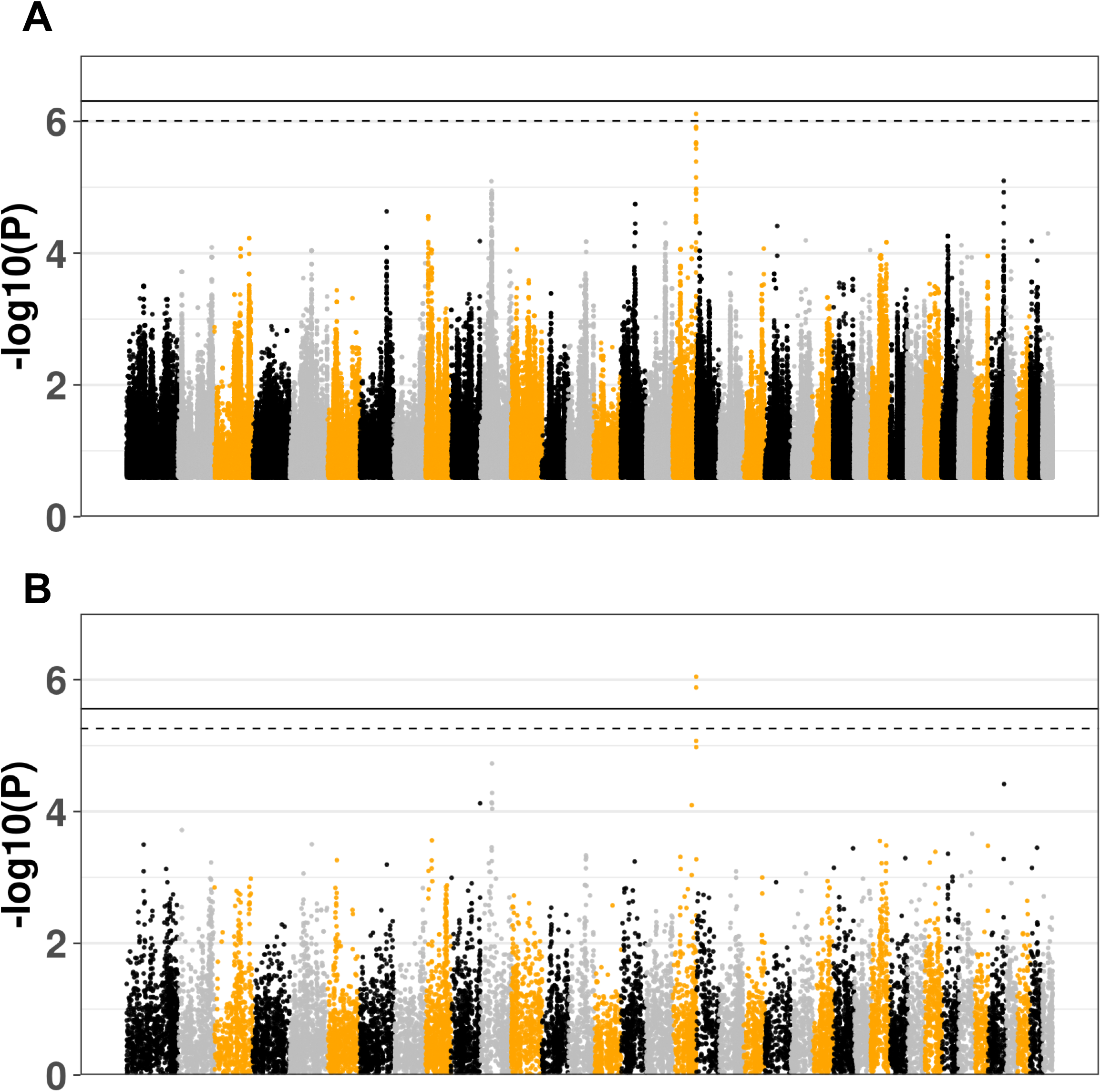
(A) SNP level analysis: Manhattan plot of the top 500K SNPs ranked by p-value. The continuous and dashed lines indicate the genome-wide and suggestive significance thresholds, respectively. (B) Gene level analysis: Manhattan plot of all the genes ranked by p-value. The continuous and dashed lines indicate the genome-wide and suggestive significance thresholds, respectively.

We obtained *λ* = 1.052, demonstrating an absence of significant population stratification after principal components adjustment (Figure S3). We did not detect any genome-wide significant variant after multiple testing correction, but one variant (rs22669389) was identified at a suggestive level of significance (adj *p* = 0.078). All of the top SNPs reported in Table 1 are located in the same region on chromosome 18 between 54,983,627 and 54,992,285 (8,658 bp) encompassing the two overlapping genes *SDHAF2* and *CPSF7*. In addition to the SNP-level-analysis, we computed a multi-marker test using the GATES method, adjusting the results for the total number of genes tested (*n* = 18,110; *p* < 2.76E-06). SNPs were assigned to a gene when ± 1,500 bp from the gene. The results showed two significant genes after Bonferroni correction: *CPSF7* (adj p = 0.016) and *SDHAF2* (adj p = 0.024). The corresponding Manhattan plot is shown in Figure 1B.

The regional plot including all of the SNPs in the region is shown in Figure 2. These results were confirmed when we considered larger distances for the SNP to gene assignments (for both ± 5,000 bp and ± 10,000 bp) (data not shown).

**Figure 2.**
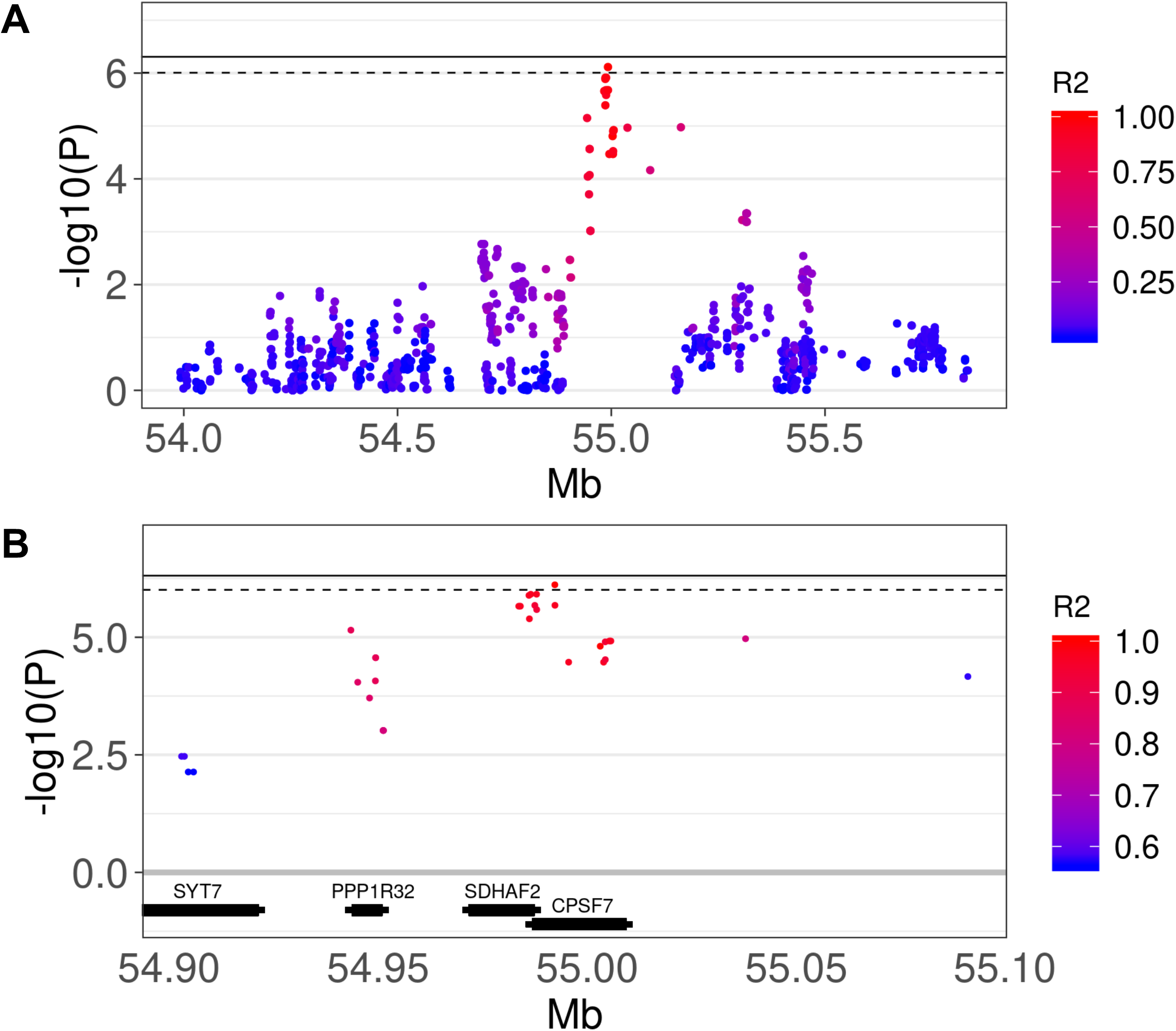
Manhattan plots showing the details of the association region in chromosome 18. The continuous and dashed lines indicate the genome-wide and suggestive significance thresholds, respectively. The color of the points indicates the Linkage Disequilibrium (expressed as R^2^) between the top (rs22669389) and the close SNPs. (A) Region ± 1 Mb from the top SNP; (B) Region ±0.1 Mb from the top SNP. Thick sections of the genes represent the actual gene region according Ensemble, the thin sections represent the surrounding regions (± 1,500).

## 4. Discussion

We were able to detect significant genetic risk factors for CIPF in the West Highland White Terrier dog breed using a GWAS including 1,839,683 informative SNPs. Applying a gene-level approach, we observed genome-wide significant signals (*p* < 2.76E-06) in *CPSF7* (Cleavage and polyadenylation specific factor 7) and *SDHAF2* (Succinate dehydrogenase complex assembly factor 2). These two overlapping genes include 15 and 8 SNPs, respectively. All of the top associated SNPs were located in introns, 5’UTR or upstream the two genes.

*CPSF7* is a human Orthologue (Gene Order Conservation Score = 100), and encodes for the 59kDa subunit of *Cleavage Factor Im*, involved in the cleavage and polyadenylation of pre-mRNAs. It is related to several mRNA process pathways, as “mRNA splicing”, “metabolism of RNA”, “mRNA 3’-end processing”, “Processing of Capped Intronless Pre-mRNA”, and “RNA Polymerase II Transcription”. In a recent study *CPSF7* was found to be involved in Lung Adenocarcinoma (LAD). Specifically, SP1 induces the promoter activity of *LINC00958* which when overexpressed drives LAD progression via the miR-625-5p/CPSF7 axis [25]. The genetic-association reported in our study might reveal the importance of CPSF7 in CIPF perhaps through the same pathologic mechanism as in lung cancer. IPF in humans is a risk factor for lung cancer, increasing the chance of development from 7% to 10% [26]. Additionally, there are several genetic, molecular and cellular mechanisms shared between lung fibrosis and lung cancer such as myofibroblast activation, endoplasmatic reticulum stress, alteration of growth factor expression, and genetic and epigenetic variations [26]. *CPFS7* has also associated with liver cancer [27].

*SDHAF2* encodes a mitochondrial protein involved in the flavination of a succinate dehydrogenase complex subunit and it has largely been associated with paragangliomas in previous literature [28,29].

In conclusion, we report for the first time the identification of genetic variants associated with CIPF in the West Highland White Terrier dog breed, located in a region encompassing the *CPFS7 and SDHAF2* genes. Our findings demonstrate some overlap with biological functions with compelling links to previously demonstrated findings in lung cancer, sharing several biological and genetic features with IPF in humans.

## Supporting information

File S1. Informed consent for biological sample collection.

Table S1. Median number of SNPs at different depth levels before quality control filtering.

Figure. S1. Scatterplot of Principal Components 1 and 2 of all of the sequenced samples before outlier removal.

Figure. S2. Distribution of the pi hat value computed between each pair of dogs included in the GWAS.

Figure. S3. QQplot showing the observed and expected distribution of GWAS p-values.

## Supplementary Materials

File S1: Informed consent for biological sample collection.

Table S1. Median number of SNPs at different depth levels before quality control filtering.

Figure. S3. QQplot showing the observed and expected distribution of GWAS p-values.

## Authors Contributions

Conceptualization, M.J.H, methodology, M.J.H, A.G.H, I.S.P and M.D.D; formal analysis, A.G.H, I.S.P, M.D.D, C.B, A.S and A.J.W; investigation M.J.H and I.S.P; data curation I.S.P. and M.D.D; writing—original draft preparation, I.S.P and M.J.H; writing—review and editing, I.S.P, M.J.H. and A.G.H; supervision, M.J.H. All authors have read and agreed to the published version of the manuscript

## References

1. Heikkilä, H.P.; Lappalainen, A.K.; Day, M.J.; Clercx, C.; Rajamäki, M.M. Clinical, bronchoscopic, histopathologic, diagnostic imaging, and arterial oxygenation findings in west highland white terriers with idiopathic pulmonary fibrosis. J. Vet. Intern. Med. 2011.

2. Clercx, C.; Fastrès, A.; Roels, E. Idiopathic pulmonary fibrosis in West Highland white terriers: An update. Vet. J. 2018.

3. Heikkilä-Laurila, H.P.; Rajamäki, M.M. Idiopathic pulmonary fibrosis in west highland white terriers. Vet. Clin. North Am. - Small Anim. Pract. 2014.

4. Daccord, C.; Maher, T.M. Recent advances in understanding idiopathic pulmonary fibrosis. F1000Research 2016.

5. Coward, W.R.; Saini, G.; Jenkins, G. The pathogenesis of idiopathic pulmonary fibrosis. Ther. Adv. Respir. Dis. 2010.

6. Lilja-Maula, L.; Syrjä, P.; Laurila, H.P.; Sutinen, E.; Palviainen, M.; Ritvos, O.; Koli, K.; Rajamäki, M.M.; Myllärniemi, M. Upregulation of alveolar levels of activin B, but not activin A, in lungs of west highland white terriers with idiopathic pulmonary fibrosis and diffuse alveolar damage. J. Comp. Pathol. 2015.

7. Krafft, E.; Lybaert, P.; Roels, E.; Laurila, H.P.; Rajamäki, M.M.; Farnir, F.; Myllärniemi, M.; Day, M.J.; Mc Entee, K.; Clercx, C. Transforming Growth Factor Beta 1 Activation, Storage, and Signaling Pathways in Idiopathic Pulmonary Fibrosis in Dogs. J. Vet. Intern. Med. 2014.

8. Krafft, E.; Laurila, H.P.; Peters, I.R.; Bureau, F.; Peeters, D.; Day, M.J.; Rajamäki, M.M.; Clercx, C. Analysis of gene expression in canine idiopathic pulmonary fibrosis. Vet. J. 2013.

9. Krafft, E.; Heikkilä, H.P.; Jespers, P.; Peeters, D.; Day, M.J.; Rajamäki, M.M.; Mc Entee, K.; Clercx, C. Serum and Bronchoalveolar Lavage Fluid Endothelin-1 Concentrations as Diagnostic Biomarkers of Canine Idiopathic Pulmonary Fibrosis. J. Vet. Intern. Med. 2011.

10. Noth, I.; Zhang, Y.; Ma, S.F.; Flores, C.; Barber, M.; Huang, Y.; Broderick, S.M.; Wade, M.S.; Hysi, P.; Scuirba, J.; et al. Genetic variants associated with idiopathic pulmonary fibrosis susceptibility and mortality: A genome-wide association study. Lancet Respir. Med. 2013.

11. Fingerlin, T.E.; Murphy, E.; Zhang, W.; Peljto, A.L.; Brown, K.K.; Steele, M.P.; Loyd, J.E.; Cosgrove, G.P.; Lynch, D.; Groshong, S.; et al. Genome-wide association study identifies multiple susceptibility loci for pulmonary fibrosis. Nat. Genet. 2013.

12. Mushiroda, T.; Wattanapokayakit, S.; Takahashi, A.; Nukiwa, T.; Kudoh, S.; Ogura, T.; Taniguchi, H.; Kubo, M.; Kamatani, N.; Nakamura, Y. A genome-wide association study identifies an association of a common variant in TERT with susceptibility to idiopathic pulmonary fibrosis. J. Med. Genet. 2008.

13. Shearin, A.L.; Ostrander, E.A. Leading the way: canine models of genomics and disease. Dis. Model. Mech. 2010.

14. Wayne, R.K.; Ostrander, E.A. Lessons learned from the dog genome. Trends Genet. 2007.

15. Lindblad-Toh, K.; Wade, C.M.; Mikkelsen, T.S.; Karlsson, E.K.; Jaffe, D.B.; Kamal, M.; Clamp, M.; Chang, J.L.; Kulbokas, E.J.; Zody, M.C.; et al. Genome sequence, comparative analysis and haplotype structure of the domestic dog. Nature 2005.

16. Ostrander, E. A.; Kruglyak, L. Unleashing the canine genome. Genome Res. 2000.

17. Gallana, M.; Utsunomiya, Y.T.; Dolf, G.; Pintor Torrecilha, R.B.; Falbo, A.K.; Jagannathan, V.; Leeb, T.; Reichler, I.; Sölkner, J.; Schelling, C. Genome-wide association study and heritability estimate for ectopic ureters in Entlebucher mountain dogs. Anim. Genet. 2018.

18. A., P.; F., B.; M.F., R.; K.W., S.; A.E., J.; K., A.; Peiravan, A.; Bertolini, F.; Rothschild, M.F.; Simpson, K.W.; et al. Genome-wide association studies of inflammatory bowel disease in German shepherd dogs. PLoS One 2018.

19. Gast, A.C.; Metzger, J.; Tipold, A.; Distl, O. Genome-wide association study for hereditary ataxia in the Parson Russell Terrier and DNA-testing for ataxia-associated mutations in the Parson and Jack Russell Terrier. BMC Vet. Res. 2016.

20. Bianchi, M.; Dahlgren, S.; Massey, J.; Dietschi, E.; Kierczak, M.; Lund-Ziener, M.; Sundberg, K.; Thoresen, S.I.; Kämpe, O.; Andersson, G.; et al. A multi-breed genome-wide association analysis for canine Hypothyroidism identifies a shared major risk locus on CFA12. PLoS One 2015.

21. Li, N.; Stephens, M. Modelling linkage disequilibrium and identifying recombination hotspots using SNP data genetics. Genetics 2003.

22. Zhou, X.; Stephens, M. Genome-wide efficient mixed-model analysis for association studies. Nat. Genet. 2012, 44, 821–4.

23. Zhou, X.; Stephens, M. Efficient multivariate linear mixed model algorithms for genome-wide association studies. Nat. Methods 2014, 11, 407–9.

24. Li, M.-X.; Gui, H.-S.; Kwan, J.S.H.; Sham, P.C. GATES: a rapid and powerful gene-based association test using extended Simes procedure. Am. J. Hum. Genet. 2011, 88, 283–93.

25. Yang, L.; Li, L.; Zhou, Z.; Liu, Y.; Sun, J.; Zhang, X.; Pan, H.; Liu, S. SP1 induced long non-coding RNA LINC00958 overexpression facilitate cell proliferation, migration and invasion in lung adenocarcinoma via mediating miR-625-5p/CPSF7 axis. Cancer Cell Int. 2020, 20, 24.

26. Ballester, B.; Milara, J.; Cortijo, J. Idiopathic Pulmonary Fibrosis and Lung Cancer: Mechanisms and Molecular Targets. Int. J. Mol. Sci. 2019, 20.

27. Fang, S.; Zhang, D.; Weng, W.; Lv, X.; Zheng, L.; Chen, M.; Fan, X.; Mao, J.; Mao, C.; Ye, Y.; et al. CPSF7 regulates liver cancer growth and metastasis by facilitating WWP2-FL and targeting the WWP2/PTEN/AKT signaling pathway. Biochim. Biophys. acta. Mol. cell Res. 2020, 1867, 118624.

28. Bausch, B.; Schiavi, F.; Ni, Y.; Welander, J.; Patocs, A.; Ngeow, J.; Wellner, U.; Malinoc, A.; Taschin, E.; Barbon, G.; et al. Clinical characterization of the pheochromocytoma and paraganglioma susceptibility genes SDHA, TMEM127, MAX, and SDHAF2 for gene-informed prevention. JAMA Oncol. 2017.

29. Smith, J.D.; Harvey, R.N.; Darr, O.A.; Prince, M.E.; Bradford, C.R.; Wolf, G.T.; Else, T.; Basura, G.J. Head and neck paragangliomas: A two-decade institutional experience and algorithm for management. Laryngoscope Investig. Otolaryngol. 2017.

