## Supplementary material for "Association of common genetic variants in the *CPSF7* and *SDHAF2* genes with Canine Idiopathic Pulmonary Fibrosis in the West Highland White Terrier": File S1. Informed consent for biological sample collection.

v 1.0

### OWNER CONSENT
### CANINE GENETICS & GENOMICS

#### Canine Saliva Collection as Essential Material for Genetic Analysis
#### Matthew J. Huentelman, PhD

I voluntarily elect to enroll my pet in a program where a sample of saliva and/or an oral cavity swab is obtained principally for DNA isolation to advance canine research. The DNA will be utilized to better understand lung disease that is prevalent in the West Highland White Terrier. Your dog's DNA sample may be collected either because your dog has been diagnosed with lung disease, because your dog is older than a certain age and has never been diagnosed with lung disease, or because your dog is from a breed that is closely related to the West Highland White.

As a voluntary participant, I have been appraised, do understand, and agree to the following.

1. I am fully aware of the risks, side effects, benefits, and consequences of collection.

2. I authorize photography for documentation (if needed), and agree to allow broad use of information gathered from my pet in scientific communications, publications, reports and/or presentations to institutional, governmental, and sponsoring agencies charged with overseeing this research program. I further understand **all personal information will be kept strictly confidential.**

3. I have been given the opportunity to ask questions regarding sample collection; all such questions have been answered to my satisfaction.

After reading the above, I voluntarily consent to my dog(s) participating in this study.

______________________________________________

**Owner (Print)**

______________________________________________

**Owner (Signature)** **Date**

**The Translational Genomics Research Institute**
**445 N. 5th Street**
**Phoenix, AZ 85004**
**(602) 343-8400**

**Owner/Agent Information**

Name:__________________________________________________

Address:________________________________________________

City:______________________State:______Zip:_______________

Phone:________________________________________________

Email:_________________________________________________

**Canine Information**

Call Name and/or Registered Name:____________________________

Reg. # (if available):______________________________________

Date of Birth:____________________________________________

Sex (Circle one): MI MN FI FS

Breed:_________________________________________________

Coat Color:_____________________________________________

Please provide any additional info you believe may be helpful (health history, etc):

**The Translational Genomics Research Institute**
**445 N. 5th Street**
**Phoenix, AZ 85004**
**(602) 343-8400**
