## Supplementary figures and images for "Association of common genetic variants in the *CPSF7* and *SDHAF2* genes with Canine Idiopathic Pulmonary Fibrosis in the West Highland White Terrier"

### Figure. S1. Scatterplot of Principal Components 1 and 2 of all of the sequenced samples before outlier removal.

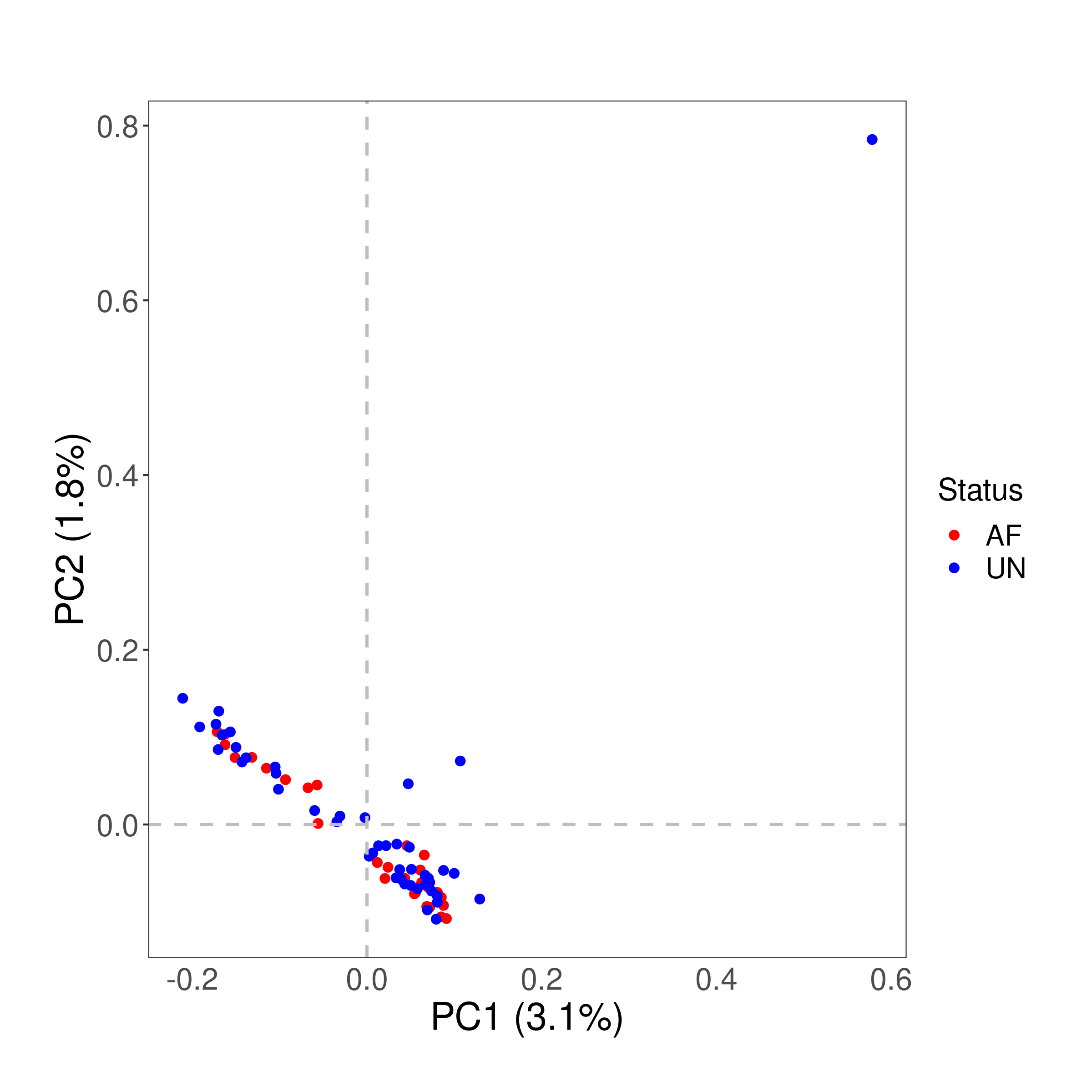

### Figure. S2. Distribution of the pi hat value computed between each pair of dogs included in the GWAS.

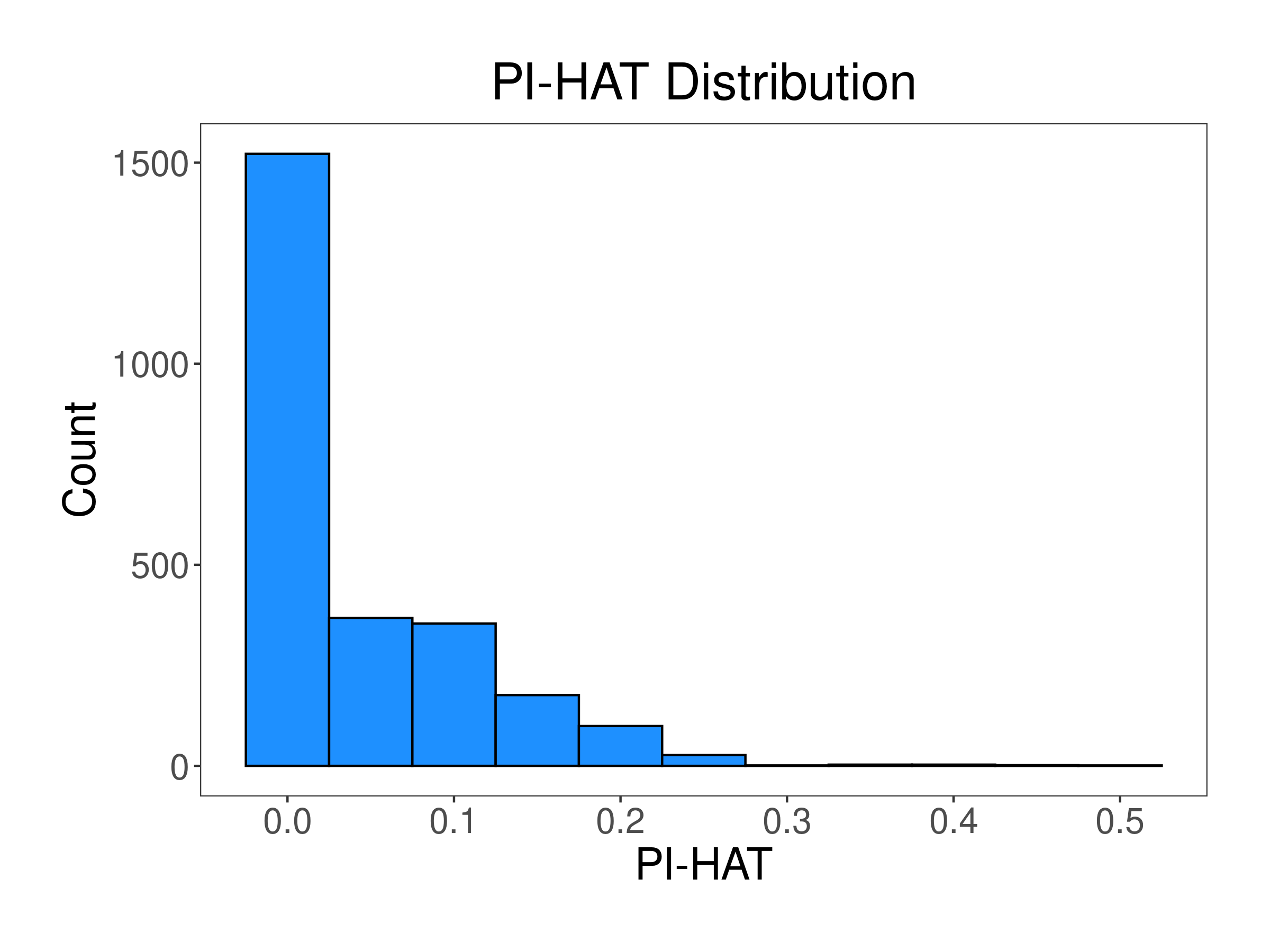

### Figure. S3. QQplot showing the observed and expected distribution of GWAS p-values.

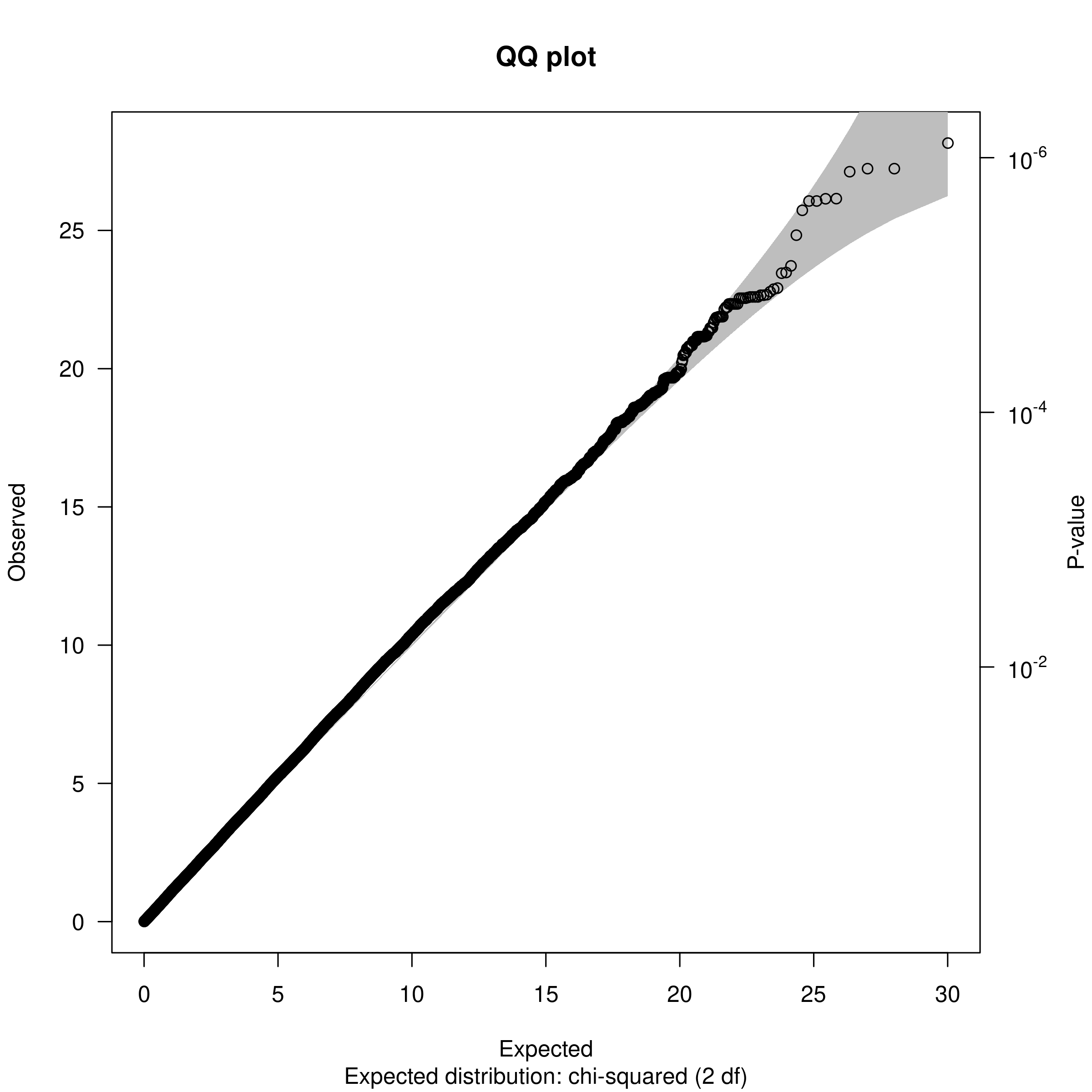
